# Conditioned Graph Reconstruction of Brain Functional Network Connectivity Reveals Interpretable Latent Axes of Sex and Fluid Intelligence

**DOI:** 10.64898/2026.02.20.707025

**Authors:** Ishaan Batta, Meenu Ajith, Vince D. Calhoun

**Affiliations:** Center for Translational Research in Neuroimaging and Data Science (TReNDS): Georgia State University, Georgia Institute of Technology, and Emory University, Atlanta, USA

**Author notes:** these authors contributed equally to this work (Ishaan Batta, Meenu Ajith).

**Keywords:** Brain Networks, Generative AI, Representation Learning, Functional Network Connectivity, fMRI, Graph Variational Autoencoder, Sex Differences, Fluid Intelligence

## Abstract

In studying the brain’s functional connectivity and its associations with clinically observed assessments, novel learning frameworks modeling its network properties in conjunction with assessment variables are crucial to uncover variable-specific patterns via meaningful encoding and reconstruction. We present a generative framework for modeling human brain functional connectivity features while retaining key network metrics and differences associated with demographic and cognitive variables. A conditional graph variational autoencoder is employed to encode static functional network connectivity (sFNC) features into a latent representation, which is then utilized for the dual purpose of reconstructing sFNC data conditioned on variables such as biological sex or fluid intelligence, and identifying discriminative connectivity features associated with the conditioning variables in the latent space. Using over 20,000 subjects from the UK Biobank, our model demonstrates high-fidelity reconstructions that preserve condition-specific network patterns, while the latent space captures interpretable patterns associated with these variables. The group differences in latent space are highlighted by one-hot probing of the latent dimensions and forward mapping to connectivity patterns. This approach provides a scalable, network-informed framework for studying brain functional connectivity and its associations with individual differences, offering potential applications in characterizing functional signatures for mental health conditions via clinically observed assessment variables.

**AUTHOR SUMMARY:** To enable the modeling of the brain functional connectivity network for encoding and reconstructing assessment-specific differences, we propose a conditional graph-based generative framework for modeling human brain functional connectivity while accounting for demographic and cognitive differences. Using a conditional graph variational autoencoder, our approach learns interpretable latent representations of functional connectivity networks derived from fMRI data. Evaluated on over 20,000 UK Biobank subjects, the model accurately reconstructs connectivity patterns outperforming baseline architectures and preserves differences associated with biological sex and fluid intelligence. By probing the latent space and mapping latent dimensions back to brain networks, we identify condition-specific connectivity features in an interpretable manner. This work provides a scalable, network-informed approach for studying individual differences in functional brain organization.

## INTRODUCTION

The brain has long been modeled as a network of regions that are functionally connected to each other. Functional magnetic resonance imaging (fMRI) is a noninvasive modality to scan full-brain functional activity that is further quantified into brain networks to study brain function and its association with clinically observed assessments. Static functional network connectivity (sFNC), usually computed as the Pearson correlation between the entire time series of pairs of brain components, offers a way to model the functional relationship between spatially separate brain areas (Du, Fu, & Calhoun, 2018).

With the advent of deep learning methods, various neural networks have been used to encode sFNC features to perform prediction (Du et al., 2018; Rahaman et al., 2023) and generative modeling tasks(L. Zhang, Wang, Zhu, Initiative, et al., 2022). Furthermore, many predictive models are often accompanied by explainability (Batta, Abrol, Calhoun, Initiative, et al., 2024) in terms of the features involved that are associated with the clinically observed target variable(s) at hand. While prediction models map neuroimaging features to clinical targets for classification or regression, generative models advance this by learning feature-to-feature mappings through a latent space, enabling the creation of synthetic features (Gong et al., 2023). By conditioning clinical variables, generative models can produce data that vary according to target values, offering both synthetic data generation and interpretable latent representations. In the case of conditioning variables, identifying sex-related differences in brain functional connectivity is clinically important as numerous neurological and psychiatric disorders exhibit sex-biased prevalence, symptomatology, and treatment response (Gualtierotti, Bressi, Garavaglia, & Brambilla, 2024). Additionally, various conditions, such as autism spectrum disorder (Napolitano et al., 2022) and Alzheimer’s disease (Fernández et al., 2024), demonstrate distinct sex-specific patterns in onset, progression, and neural correlates. Alongside sex, cognitive traits such as fluid intelligence also play a crucial role in brain function and are linked to variability in neural connectivity patterns (Qiu et al., 2025). Therefore, in modeling functional connectivity, it is essential to develop approaches that incorporate sFNC network structure and capture meaningful latent representations associated with key conditioning variables (Sidulova & Park, 2023). Moreover, approaches based on graph neural networks capturing non-linear relations in graph structures can summarize non-linear relations in the data at multiple scales, which is harder to achieve with standard machine learning models.

sFNC features can be utilized to create graphs for analyzing their properties and perform predictions as a functional connectome (Bassett & Sporns, 2017). For modeling the brain as a network using functional connectivity, graph neural networks (GNNs) provide an effective framework to encode the multi-level functional associations between brain areas within a network paradigm (Bessadok, Mahjoub, & Rekik, 2022; Mohammadi & Karwowski, 2024). GNNs have found use in many applications, ranging from brain state prediction (Lu & Uddin, 2023), disease classification (Mohanraj, Raman, & Ramanathan, 2024), and predicting longitudinal trajectories (Bessadok et al., 2022). In terms of successful generative modeling, variational autoencoders (VAEs) have been extensively used in neuroimaging applications for clustering patterns of brain activity and detecting subtle associated feature changes (Qiang et al., 2020). Moreover, the learned lower-dimensional latent features in the VAE serve as an effective way of compressing complex features for reconstruction and understanding underlying mechanisms of feature generation and associations (Zhao, Adeli, Honnorat, Leng, & Pohl, 2019).

Machine learning models have emerged as powerful tools in functional neuroimaging, particularly for synthesizing and analyzing sFNC derived from rs-fMRI data (Rahaman et al., 2023). Prior studies have demonstrated the utility of deep generative models such as VAEs (Y. Zhang, Liu, Zhang, & Dunson, 2024), generative adversarial networks (GANs) (Qiang et al., 2023), and multimodal transformer models (Abrol et al., 2021; Bi, Abrol, Jia, Sui, & Calhoun, 2024) in capturing complex brain connectivity patterns. However, most existing generative models for fMRI and sFNC data lack integration of subject-level covariates, limiting their ability to generate biologically and cognitively meaningful variations in synthetic data (Ajith et al., 2024; Lewis, Miller, Gazula, & Calhoun, 2023). In most cases, these models are evaluated only on relatively small, homogeneous datasets. Moreover, they also lack interpretable latent representations (J.-H. Kim et al., 2021) that can be meaningfully linked back to brain network patterns used to explore condition-specific differences in connectivity. Hence, we introduce the Conditional Graph Variational Autoencoder (C-GVAE), which conditions on subject-specific covariates while learning robust latent representations of brain networks. Unlike earlier methods, this model incorporates GATv2 (Brody, Alon, & Yahav, 2021) encoders to extract meaningful latent representations from graph-structured sFNC inputs, which are then conditioned on either biological sex or fluid intelligence scores before being decoded through a multilayer perceptron (MLP) into synthetic connectivity graphs. We test our framework to show that the resconstructed sFNCs enhance predictive power while also retaining key network properties (assortativity, clustering coefficient, etc.) from the real data. The proposed framework offers four primary contributions: (1) it enables conditional generation of high-fidelity sFNC data in the form of brain connectivity networks that captures variability associated with both sex and fluid intelligence, validated using two large sub-cohorts from the UK Biobank and further tested on the HCP dataset; (2) it preserves meaningful group-level differences in brain functional connectivity patterns based on each conditioning variable and enhances classification performance in sex and fluid intelligence prediction tasks; (3) it preserves the key local and global graph properties from the real data such that the generated sFNC graphs are not structurally different; and (4) it learns interpretable latent representations, from which key discriminative dimensions can be visualized and decoded into brain networks. Overall, our framework is successfully able to achieve the multifaceted goal of synthesizing realistic functional connectivity features, retaining key discretionary patterns for the target conditional variables, preserving key network metrics, and learning meaningful latent representations associated with these differences.

## METHODS

### Dataset and Preprocessing

The neuroimaging dataset used in this study was from the UK Biobank database (Miller et al., 2016) and included 20,000 participants aged between 55 and 88 years (mean age 71.26 ± 7.44). The cohort consisted of 9,393 females (46.96%) and 10,607 males (53.04%). In this analysis, two separate conditioning variables (assessment measures) were used: sex and fluid intelligence score. Fluid intelligence is a standardized cognitive assessment that measures reasoning and problem-solving abilities, independent of acquired knowledge. The fluid intelligence scores in this cohort ranged from 0.0 to 13.0 (mean score of 6.11 ± 7.04). Additionally, to evaluate the generalizability of the findings, the Human Connectome Project (HCP) dataset (Van Essen et al., 2013) was employed as an external validation cohort.

In case of neuroimaging data, all scans were performed using a 3-Tesla Siemens Skyra scanner equipped with a 32-channel head coil. Rs-fMRI data were acquired using a gradient-echo echo planar imaging (GE-EPI) sequence. Preprocessing involved motion correction, normalization, high-pass temporal filtering, and geometric distortion correction. Additional steps included EPI unwarping and gradient distortion correction. Noise and artifacts were removed using Independent Component Analysis (ICA), a data-driven method that separates mixed signals into statistically independent components, combined with FMRIB’s ICA-based X-noiseifier (FIX). Finally, the data were registered to the Montreal Neurological Institute (MNI), a standardized brain coordinate system, using an EPI template and smoothed with a 6×6×6 mm full width at half maximum Gaussian kernel.

After preprocessing, a fully automated spatially constrained ICA was applied to the rs-fMRI data (Du et al., 2020). We used the Neuromark fMRI 1.0 template, consisting of 53 reproducible, data-driven independent components (ICs) representing well-characterized functional brain networks. These components are organized into seven functional subdomains: Subcortical (SC), Auditory (AUD), Sensorimotor (SM), Visual (VIS), Cognitive Control (CC), Default Mode Network (DMN), and Cerebellar (CB). Component definitions and subdomain assignments are summarized in Table 1.

**Table 1:**
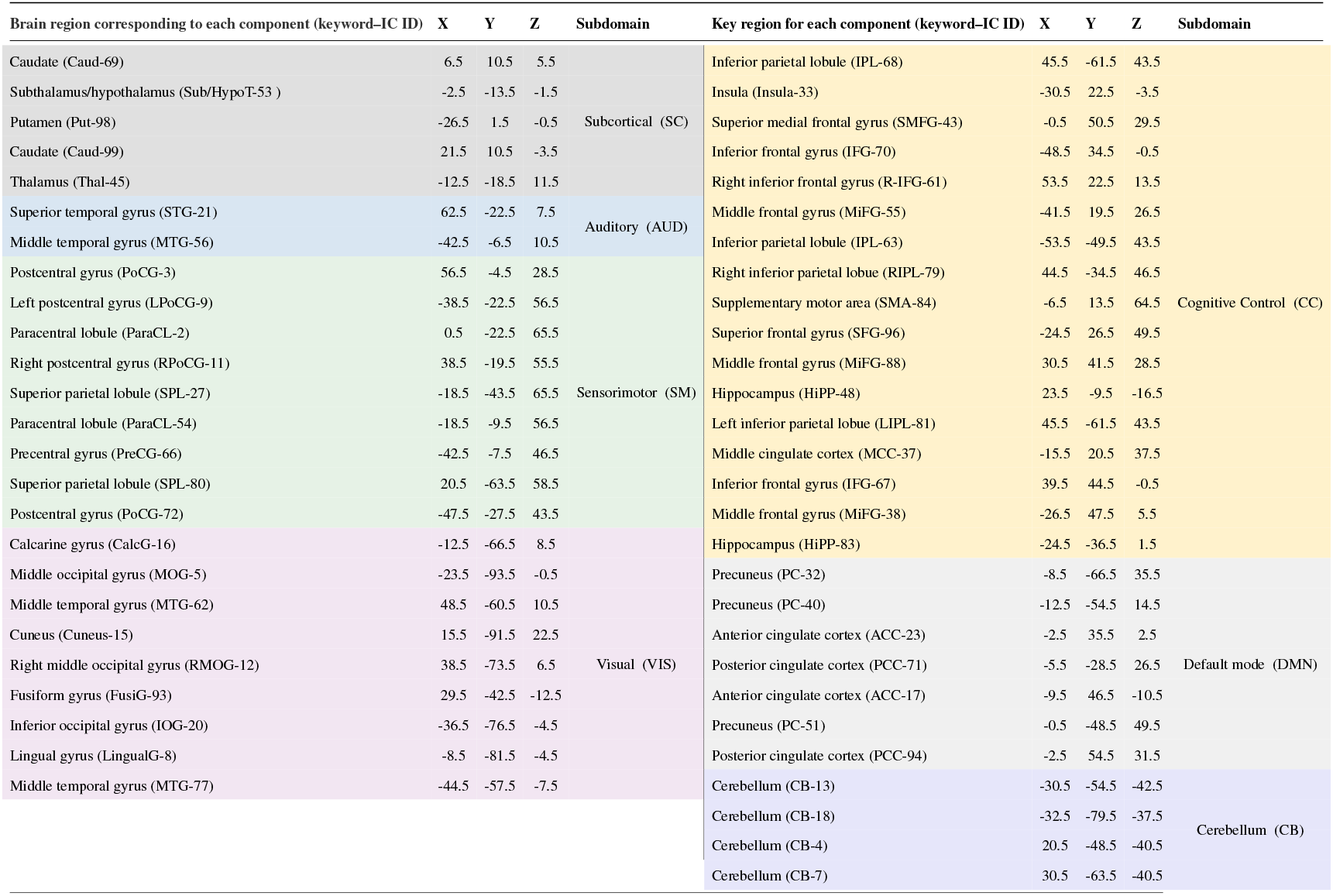
The table lists the peak MNI coordinates and associated brain regions for 53 spatially constrained ICA components derived from the Neuromark template for fMRI data. These ICs are grouped into seven functional subdomains, each highlighted in a distinct color, and available in the GIFT toolbox. Descriptive keywords in parentheses are used to reference components in subsequent figures. The components were also used to compute 53 × 53 static functional connectivity matrices for input to our model.

Static functional network connectivity (sFNC) was computed by estimating Pearson correlation between all pairs of IC time courses, yielding 53 × 53 connectivity matrices for each subject. These Ics and subdomains are used throughout Figure 2, Figure 3, and Figure 6 to visualize connectivity patterns, group-level pairwise statistics, and network interactions using connectivity matrices and connectograms.

### Conditional Graph Variational Autoencoder (C-GVAE)

We propose a Conditional Graph Variational Autoencoder (C-GVAE) for generating synthetic sFNC matrices derived from rs-fMRI data (Figure 1). The model extends the standard VAE framework to handle graph-structured data with conditioning on auxiliary attributes such as sex. The input to the model consists of subject-wise graphs that are constructed from sFNC matrices. To reduce noise and enforce sparsity, a thresholding procedure is applied. Specifically, all diagonal elements are set to zero to eliminate self-connections. Then, for each off-diagonal element, the original signed value is retained only if its absolute magnitude exceeds a fixed threshold *τ* = 0.1; otherwise, the element is set to zero. The threshold *τ* was selected based on a density-based criterion to ensure a consistent and interpretable level of sparsity in the graphs, resulting in sparse, signed, weighted adjacency matrices while preserving the strongest functional connections. The same threshold value is applied to all subjects and datasets, resulting in graphs with comparable density. As *τ* is fixed a priori and not optimized using outcome variables or validation performance, the thresholding procedure does not introduce information leakage between training and evaluation data.

**Figure 1:**
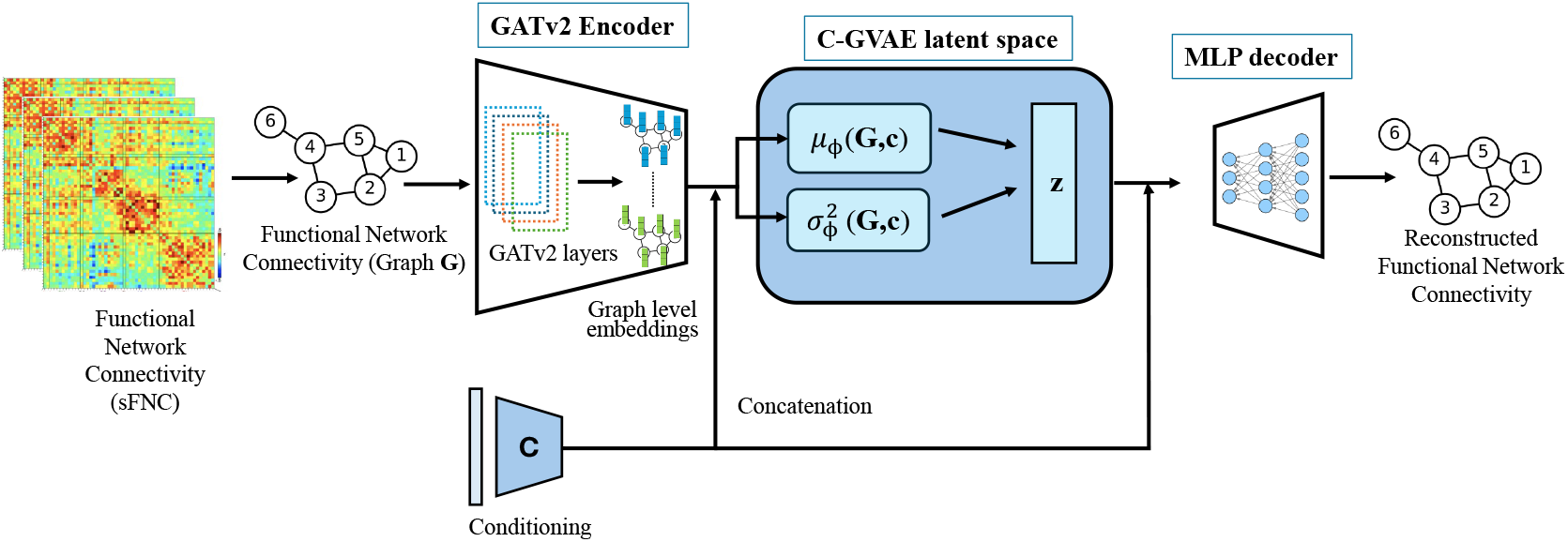
Conditional variational autoencoder (C-GVAE) with a GATv2 encoder was used for modeling functional connectivity networks. The encoder maps the sFNC matrix to a graph embedding, which is concatenated with the conditioning variable (sex) to compute the latent distribution parameters ***µ***_*ϕ*_(𝒢, **c**) and 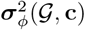. A latent sample is reparameterized, concatenated again with the conditioning variable, and passed to the MLP decoder to reconstruct the sFNC graph. The model is trained end-to-end to generate conditioned sFNC graphs.

Each input sample is represented as a graph 𝒢 = (𝒱, ℰ), where nodes correspond to ICs and edges represent significant functional associations between them. Each node *v* ∈ 𝒱 is associated with a feature vector **h**_*v*_ ∈ ℝ^*N*^, forming the node feature matrix **X** ∈ ℝ^*N*×*N*^, where the features correspond to the rows of the thresholded sFNC matrix. The graph connectivity is captured by the edge index, which includes all non-zero off-diagonal entries of the thresholded adjacency matrix. Meanwhile, sex information is encoded as a binary float tensor to be used as a conditioning variable in the latent space of C-GVAE.

#### Graph Encoder Variants

To encode the graph-structured sFNC data into a latent space suitable for generative modeling, several GNN architectures were employed as the encoder component of the C-GVAE framework. GNNs are a class of deep learning models specifically designed to process non-Euclidean data, such as graphs, by jointly leveraging node attributes and connectivity patterns among nodes (Bronstein, Bruna, LeCun, Szlam, & Vandergheynst, 2017; Wu et al., 2020). Unlike traditional neural networks that assume images or sequences, GNNs operate by iteratively aggregating and transforming information from a node’s local neighborhood (Wu et al., 2020). This enables the learning of expressive node-level and graph-level representations (Jiang, Zhang, Lin, Tang, & Luo, 2019). Through message passing and attention-based mechanisms, GNNs model complex relational patterns, making them particularly well-suited for applications such as brain network analysis (B.-H. Kim & Ye, 2020; Li et al., 2021).

The first encoder architecture evaluated was the Graph Convolutional Network (GCN)(B.-H. Kim & Ye, 2020), which updates each node’s features by aggregating information from its local neighborhood using a normalized adjacency matrix. This operation preserves local graph structure and incorporates contextual information from connected nodes. Building on this, Graph Attention Networks (GATs)(Veličković et al., 2017) introduce an attention mechanism that learns the relative importance of neighboring nodes during feature aggregation. Attention coefficients are computed between a node and its immediate neighbors and used to perform a weighted sum of neighboring features, enabling the model to focus on the most informative parts of the neighborhood. The Graph Isomorphism Network (GIN)(Xu, Hu, Leskovec, & Jegelka, 2018) was also implemented, which uses a sum-based neighborhood aggregation followed by an MLP. This design enables the model to capture fine-grained structural differences between graphs.

Finally, the Graph Attention Network v2 (GATv2) (Brody et al., 2021) improves upon the GAT model by addressing its limitations. GATv2 offers richer representation learning through dynamic attention, enabling better modeling of functional brain network structures. In the context of graph representation learning for sFNC data, the superior performance of the GATv2 encoder stems from its expressive attention mechanism. The node features are processed through *L* layers of GATv2 convolutions. Each layer computes attention scores between a node and its neighbors as follows:

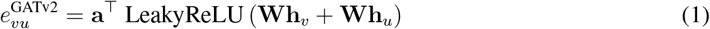

where **W** is a learnable linear transformation, **a** is a learnable projection vector, and the attention coefficients are normalized via a softmax over the neighborhood of each node:

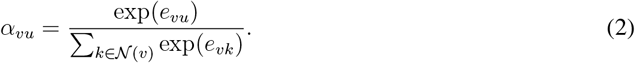

The outputs of multi-head attention are concatenated and passed through batch normalization, a LeakyReLU activation, and dropout for regularization. Finally, a global additive pooling operation aggregates node-level representations into a graph-level embedding, which is mapped through fully connected layers to produce the latent vector **z** used in the proposed model.

The effectiveness of this representation learning is closely tied to the underlying attention formulation. In contrast to the original GAT, GATv2 computes attention scores using a nonlinear combination of node features. By comparison, GAT computes attention coefficients as follows:

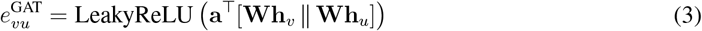

In GAT, the attention mechanism relies on a fixed linear projection of concatenated node features, limiting its flexibility, whereas GATv2 introduces a nonlinear combination of node features prior to projection using a learnable linear transformation. This enables the model to capture more complex, learnable interactions between node pairs compared to the fixed concatenation in GAT.

As a result, the normalized attention weights in GATv2 assign greater importance to informative neighbors based on both local and global context. This is particularly important in modeling brain connectivity graphs, where not all regions contribute equally to a cognitive function or disorder. Thus, the architectural advantages of GATv2 directly translate to improved performance in learning and encoding functional brain networks (Hu, Cao, Li, Dong, & Li, 2021; Safai et al., 2022).

#### Latent Space Conditioning and Generation

Let **c** denote the conditioning variable, which is the binary sex label (male or female). Conditioning is applied at both the encoder and decoder stages of the proposed C-GVAE model. Given an input graph 𝒢, the encoder first generates a graph-level representation, which is then concatenated with the conditioning variable **c**. This concatenated representation is used to parameterize a conditional approximate posterior distribution *q*_*ϕ*_(**z** | 𝒢, **c**), where **z** ∈ ℝ^*k*^ denotes the latent variable. Consequently, the encoder outputs the mean ***µ***_*ϕ*_(𝒢, **c**) and variance 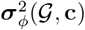 of a diagonal Gaussian distribution. The latent sample **z** is obtained via the reparameterization trick and concatenated with the same conditioning variable **c**. Following this, the decoder is implemented as a multilayer perceptron that takes a sampled latent vector **z** concatenated with the conditioning variable **c**. Finally, it reconstructs the input sFNC matrix by modeling *p*_*θ*_(𝒢 | **z, c**).

A standard Gaussian prior *p*(**z**) = 𝒩(**0, I**) is assumed over the latent space and is shared across all conditions. Conditioning is therefore introduced exclusively through the encoder and decoder, rather than through a condition-specific prior. Therefore, the proposed C-GVAE model is trained by maximizing the conditional evidence lower bound (ELBO).

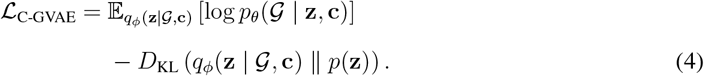

Additionally, the total training objective combines a reconstruction loss and a Kullback–Leibler divergence term with respect to a shared standard Gaussian prior:

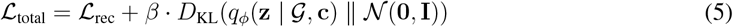

where *β* is a weighting coefficient to control the KL term. This conditional formulation enables the generation of synthetic sFNC matrices conditioned on subject-level attributes, facilitating the analysis of group-specific patterns in functional brain connectivity.

#### Experimental Setup

The proposed C-GVAE model was utilized to generate synthetic sFNC data, with a specific focus on incorporating sex and fluid intelligence conditioning information into the model. To assess its generalizability and mitigate the risk of overfitting, the model was trained and evaluated on two independent subsets of the UK Biobank dataset, referred to as UKB1 and UKB2, each containing 10,000 subjects. Additionally, to further validate cross-dataset robustness, the model was tested on the HCP dataset comprising 833 subjects. The architecture of the C-GVAE comprised a graph-based encoder specifically designed to process graph-structured data. This encoder used GATv2 to extract meaningful graph representations from weighted graphs, while an MLP served as the decoder, reconstructing graphs from latent embeddings.

The training process began by partitioning the dataset into a fixed 10% external test set, with the remaining 90% used for training and validation. To ensure robustness and prevent overfitting, a random subsampling approach with *N* = 10 splits was employed. This cross-validation strategy helped improve the model’s generalizability. The C-GVAE model was trained over 100 epochs using the Adam optimizer with weight decay and a learning rate scheduler, optimizing a composite loss function that combined both reconstruction loss and KL divergence. For each split, the model with the lowest validation loss was selected and saved. Once trained, the encoder projected the graphs into a latent space, which was then concatenated with the conditioned labels and passed as input to the decoder MLP.

Model performance was assessed on the held-out test set using several evaluation metrics, including MSE, Frobenius norm, correlation, and graph-level topological statistics. These metrics were used to evaluate the fidelity and structural quality of the reconstructed graphs. For hyperparameter optimization of the C-GVAE model, we employed Optuna (Akiba, Sano, Yanase, Ohta, & Koyama, 2019), a framework that automates the process of finding optimal hyperparameters through an efficient probabilistic model. The final model configuration included a batch size of 16, a GATv2 encoder hidden dimension of 128, and an MLP decoder hidden dimension of 128, with a latent space dimensionality of 50. The encoder and decoder architectures consisted of four and two layers, respectively. The model was trained at a learning rate of 0.0001 and could accommodate graphs with up to 53 nodes. In terms of computational complexity, the overall training time was 21 minutes and 40 seconds, and the test time per sample was 1.01 ms. Additionally, the model contained approximately 1.90 million parameters, and all experiments were conducted on a single NVIDIA A40 GPU with 4 GB of allocated RAM.

#### Model selection and ablation study for GraphVAE

For model selection within the GraphVAE framework, five different encoder architectures were systematically evaluated in independent sets of UK Biobank datasets to determine the most effective approach for graph representation learning. The different encoders included the GIN, GCN, GAT, GATv2, and a fully connected MLP baseline. During this comparison, all models shared a fixed MLP decoder. Each configuration was trained and validated using consistent data splits and hyperparameters. Performance was assessed based on correlation, MSE, and Frobenius norm, as shown in Table. 2. Among all architectures, GAT and GATv2 demonstrated the best overall performance and captured both local and global graph structures more effectively. Given its slightly superior results in both datasets, GATv2 was selected as the encoder for the final C-GVAE model.

**Table 2:** Performance comparison of different encoder architectures within the C-GVAE framework.

| Dataset | GraphVAE | Evaluation metrics |  |  |
| --- | --- | --- | --- | --- |
| UKB1 | Encoder | Correlation | MSE | Forbenius norm |
|  | MLP | 0.7890+/-0.0008 | 0.0573+/-0.0002 | 12.5372+/-0.0246 |
|  | GCN | 0.8387+/-0.0006 | 0.0445+/-0.0002 | 11.0691+/-0.0226 |
|  | GIN | 0.8478+/-0.0006 | 0.0418+/-0.0002 | 10.7332+/-0.0195 |
|  | GAT | 0.8859+/-0.0004 | 0.0315+/-0.0001 | 9.3315+/-0.0158 |
|  | **GATv2** | **0.8878+/-0.0008** | **0.0310+/-0.0002** | **9.2519+/-0.0290** |
| UKB2 | MLP | 0.7777+/-0.0006 | 0.0439+/-0.0001 | 10.9914+/-0.0164 |
|  | GCN | 0.8275+/-0.0004 | 0.0347+/-0.0001 | 9.7675+/-0.0135 |
|  | GIN | 0.8379+/-0.0007 | 0.0325+/-0.0001 | 9.4672+/-0.0200 |
|  | GAT | 0.8797+/-0.0006 | 0.0246+/-0.0001 | 8.2297+/-0.0192 |
|  | **GATv2** | **0.8808+/-0.0008** | **0.0241+/-0.0001** | **8.1478+/-0.0245** |

We also conducted an ablation study to analyze the impact of incorporating conditioning variables into the GraphVAE framework. As shown in Table 3, conditioning the model on sex and fluid intelligence consistently improved performance across both UKB1 and UKB2 subsets compared to the unconditional baseline. Specifically, sex conditioning resulted in the lowest MSE and Frobenius norm values, indicating enhanced reconstruction accuracy and structural fidelity in the reconstructed graphs. While fluid intelligence conditioning also provided performance gains, the improvements were slightly less pronounced than those observed with sex. These results underscore the benefits of incorporating subject-specific covariates to guide the generative process. Additionally, the proposed CGVAE model was also compared to the standard Conditional VAE (CVAE) for both sex and fluid intelligence conditioning on the UKB1 and UKB2 datasets. For sex conditioning, the CVAE achieved average Frobenius norms of 11.0696 ± 0.0196 (UKB1) and 11.1373 ± 0.0156 (UKB2), MSE values of 0.0445 ± 0.0002 and 0.0451 ± 0.0001, and correlation scores of 0.7791 ± 0.0008 and 0.7783 ± 0.0007, respectively. For fluid intelligence, the CVAE yielded Frobenius norms of 10.8861 ± 0.0094 and 10.8614 ± 0.0153, MSE values of 0.0430 ± 0.0001 and 0.0428 ± 0.0001, and correlations of 0.7836 ± 0.0005 and 0.7847 ± 0.0007 on UKB1 and UKB2. Hence, in comparison, the CGVAE model (see Table 3) consistently outperforms the CVAE across all metrics and conditioning variables, demonstrating lower reconstruction errors and higher correlation scores.

**Table 3:** Ablation study showing model performance across different conditioning settings (unconditional, sex as conditioning, and fluid intelligence as conditioning) for two UK Biobank subsets (UKB1 and UKB2).

| Condition type | UKB1 |  |  | UKB2 |  |  |
| --- | --- | --- | --- | --- | --- | --- |
|  | Correlation | MSE | Frobenius norm | Correlation | MSE | Frobenius norm |
| Unconditional | 0.8878±0.0008 | 0.0310±0.0002 | 9.2519±0.0290 | 0.8808±0.0008 | 0.0241±0.0001 | 8.1478±0.0245 |
| Sex | 0.8866±0.0004 | 0.0266±0.0001 | 7.9019±0.0151 | 0.8840±0.0006 | 0.0228±0.0001 | 7.9415±0.0204 |
| Fluid intelligence | 0.8824±0.0005 | 0.0238±0.0001 | 8.0956±0.0162 | 0.8808±0.0005 | 0.0241±0.0001 | 8.1476±0.0174 |

Finally, to further evaluate the robustness and generalizability of the proposed model, the unconditional GraphVAE was tested on the independent HCP dataset. It achieved a Frobenius norm of 7.7025 ± 0.0383, an MSE of 0.0217 ± 0.0002, and a correlation of 0.8600 ± 0.0014. This demonstrates that the model retains strong performance when applied to datasets beyond the UK Biobank.

#### Reconstructed Data Characteristics and Group Differences

After selecting the best-performing GATv2 model, we studied the properties of the reconstructed sFNC data based on independent training on UKB1 and UKB2 datasets. Figure 2 shows the sFNC matrices from three randomly selected subjects from UKB1 and UKB2, along with the mean and standard deviation of the sFNC values for each dataset. It can be noticed that the mean sFNC values for reconstructed and real data are very similar to each other. A two-sample t-test with FDR correction was performed to compare the feature-wise sample means for the sFNC values. While some entries (37 functional connections) did survive the FDR correction procedure, the actual differences in the mean values between reconstructed and real data for these entries were very minimal (*<* 0.05), thus indicating that our GraphVAE framework is able to generate realistic sFNC data. Additionally, we also looked into the group differences between the male and female sFNC features independently for both reconstructed as well as real data. Figure 3 shows the results from the two-sample t-tests with FDR correction to compare sFNC features from male and female groups, performed independently for real and reconstructed data from UKB1 and UKB2 datasets. As can be noticed from Figure 3, the sFNC features with significant sex differences are very well preserved in the reconstructed data, with a very low false positive rate of only 0.015 for UKB1 and 0.014 for UKB2. For fluid intelligence scores, we performed Pearson correlation analysis with each element of the SFNC matrix for the data, followed by FDR correction. The results, visualized in Figure 3 signify a similar pattern where reconstructed data very well preserves significant associations with fluid intelligence scores.

**Figure 2:**
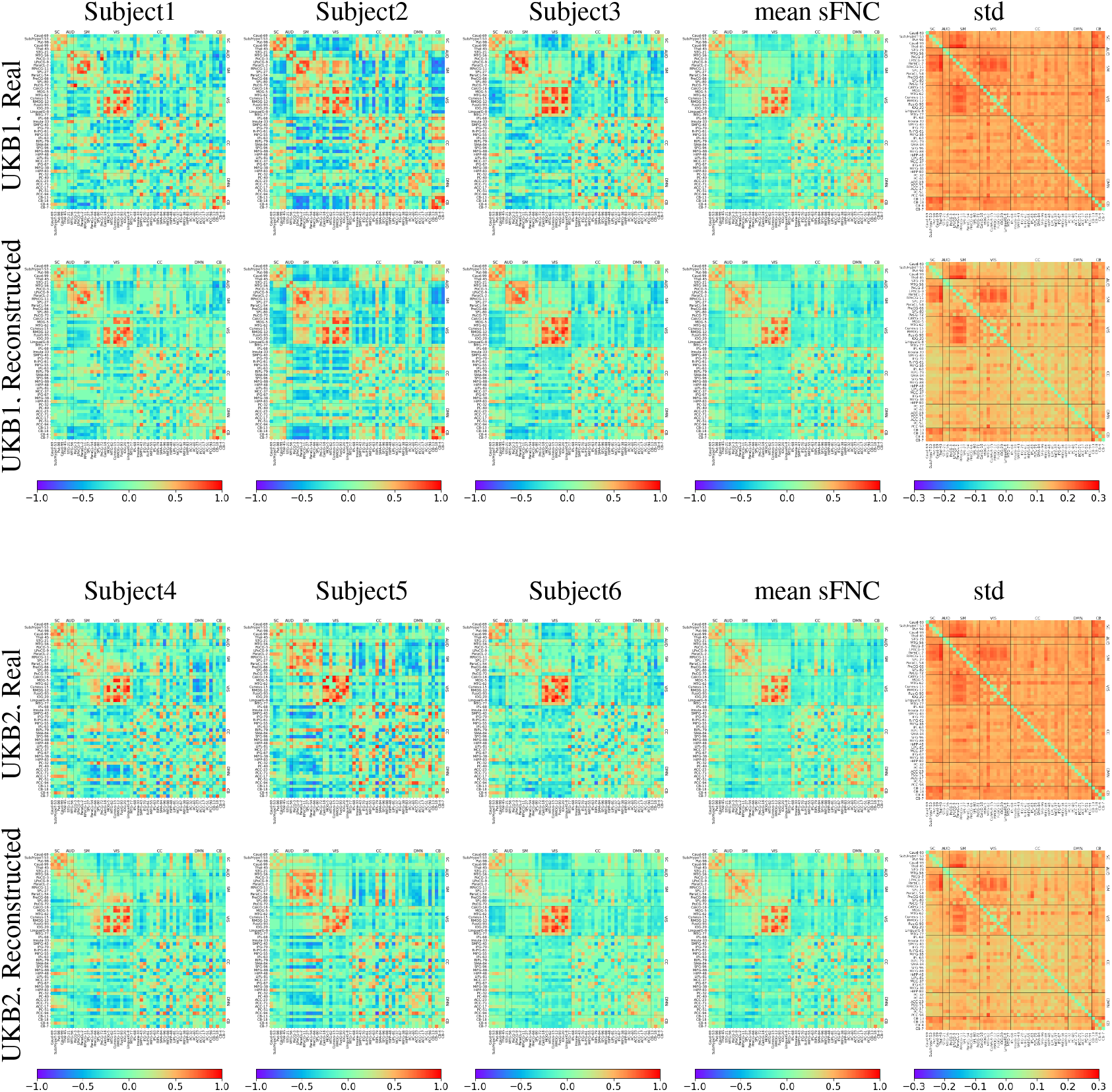
Static functional network connectivity (sFNC) computed for a subject as the Pearson correlation between component time series for 53 Neuromark brain components, divided into 7 functional subdomains. The figure shows real and reconstructed sFNCs for three random subjects each, along with element-wise mean and standard deviation across the dataset for the UKB1 and UKB2 datasets.

**Figure 3:**
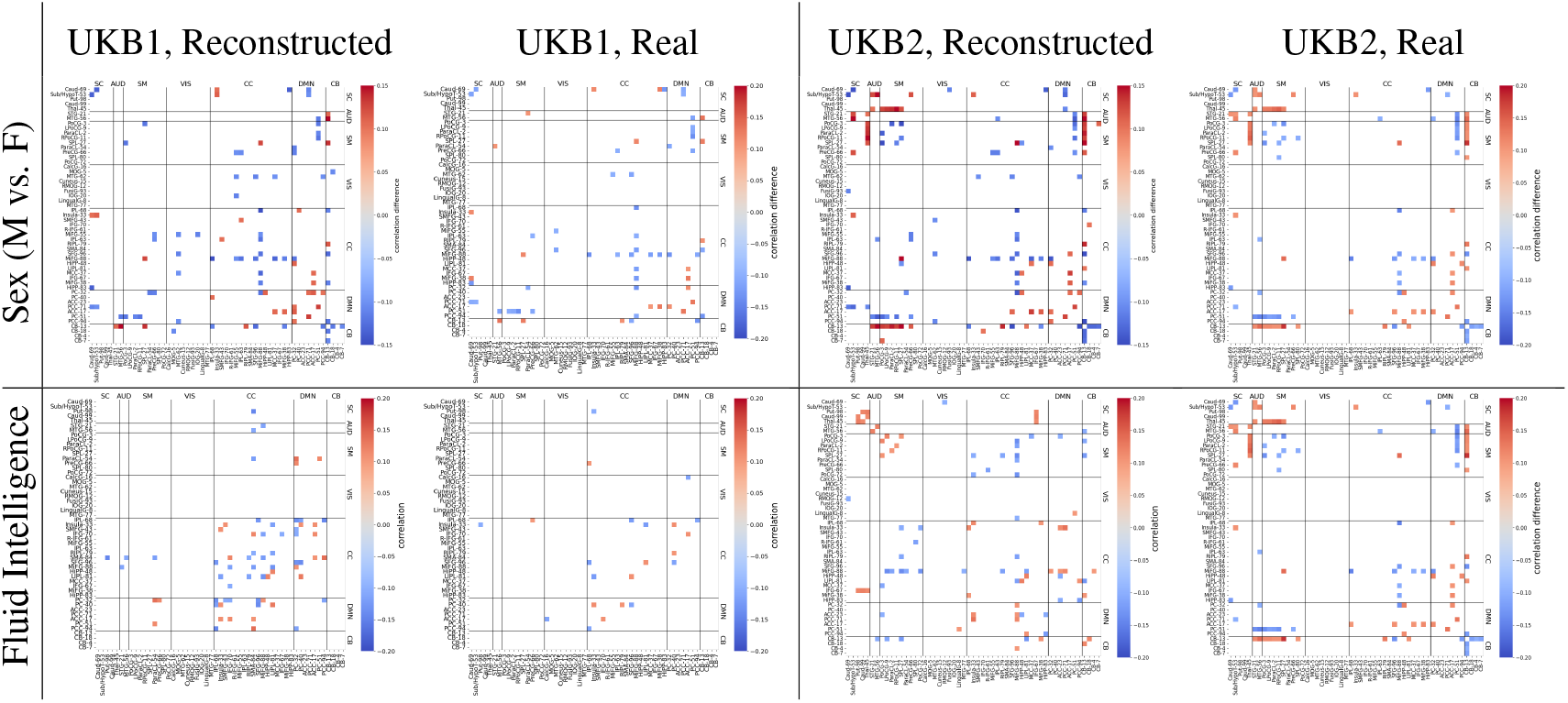
Differences between means of male and female SFNC features based on a two-sample t-test with FDR correction (top row) are visualized values (for entries with *p <* 0.01 after FDR correction). For fluid intelligence, correlation between each SFNC feature and the fluid intelligence score is shown for significantly non-zero entries based on two-sided t-test after FDR correction. Results are shown for both real and reconstructed data for UKB1 and UKB2 datasets.

#### Predictive capabilities of reconstructed data

To check for predictive properties of the reconstructed data, we trained a support vector machine (SVM) based classifier for cross-dataset sex classification (M vs F) between the reconstructed and real data. A total of three scenarios were studied independently. For the first scenario (*X*_*te*_ → *X*_*te*_), we did both training and testing using the SVM classifier on real held-out test data with 1000 subjects. For the second scenario (*X*_*te*_ → *X*_*gen*_), we trained the classifier on real held-out test data not used during the C-GVAE training, and tested it on the data reconstructed by C-GVAE. For the third scenario (*X*_*gen*_ → *X*_*te*_), the classifier training was done on the reconstructed data, and the testing was done on the real held-out test data. For all three scenarios and both datasets (UKB1 and UKB2), the SVM classifiers were trained independently with 5-fold cross-validation to optimize for the parameters using the radial basis function (RBF) kernel. The same analysis setup was performed for fluid intelligence score, but for a regression task using support vector regression (SVR).

Table 4 summarizes the performance results from these three scenarios for both datasets. It can be noted that the while the *X*_*te*_ → *X*_*te*_ scenario (training and testing within real data) performs worse than the *X*_*te*_ → *X*_*gen*_ scenario (training on real and testing on reconstructed data). This could be because the reconstructed data encodes the sex differences and fluid intelligence characteristics more significantly (as seen in Figure 3) than the real data, thus leading to a better separability for classification and fit for regression even when being used as a testing dataset. On the other hand, the *X*_*gen*_ → *X*_*te*_ (training on reconstructed and testing on real data) scenario outperforms all other scenarios, which further shows that the C-GVAE with conditioning generates data with patterns that not just retain but also enhance sex differences and fluid intelligence trends.

**Table 4:** Predictive performance comparison for cross-dataset sex classification with SVM for Sex and SVR for fluid intelligence using data reconstructed by C-GVAE (*X*_*gen*_) and unseen test set (*X*_*te*_) from real data for UKB1 and UKB2. *A* → *B* represents training the classifier using data *A* and testing it on *B*.

| Data | Scenario | Sex |  |  |  | Fluid Intelligence |  |  |
| --- | --- | --- | --- | --- | --- | --- | --- | --- |
|  |  | Accuracy | Precision | Recall | ROC AUC | MSE | R2 | R |
| UKB1 | $X_{te} \rightarrow X_{te}$ | 0.810 | 0.806 | 0.765 | 0.875 | 0.0506 | 0.0235 | 0.2825 |
| | $X_{te} \rightarrow X_{gen}$ | 0.848 | 0.826 | 0.841 | 0.932 | 0.0375 | 0.0776 | 0.3860 |
| | $X_{gen} \rightarrow X_{te}$ | 0.946 | 0.976 | 0.903 | 0.992 | 0.0725 | -0.7860 | 0.4075 |
| UKB2 | $X_{te} \rightarrow X_{te}$ | 0.790 | 0.820 | 0.709 | 0.874 | 0.0410 | 0.0076 | 0.2325 |
| | $X_{te} \rightarrow X_{gen}$ | 0.851 | 0.816 | 0.883 | 0.926 | 0.0360 | 0.0743 | 0.3636 |
| | $X_{gen} \rightarrow X_{te}$ | 0.953 | 0.973 | 0.926 | 0.994 | 0.0379 | 0.0242 | 0.4486 |

#### Assessment of reconstructed network metrics

To quantitatively assess whether the proposed C-GVAE preserves meaningful graph-level topology beyond edge-wise reconstruction, a comprehensive set of network properties was compared between the ground-truth and reconstructed sFNC graphs. A collection of 15 graph property statistics was computed on weighted sFNC graphs to capture both local and global network organization. The basic structural properties included the number of nodes, the number of edges, and graph density. These measures summarize overall network size and sparsity. Local connectivity in the graphs was characterized by using the weighted node degree. The core nodes and overall integration were captured by extracting maximum, minimum, and average values. Topological organization was further assessed using weighted degree assortativity. This measure reflects the tendency of nodes to connect to others with similar connectivity strength. Clustering measures included the average weighted clustering coefficient and global clustering. They quantify local segregation and overall triangle density. Additional triangle-based statistics included the total number of triangles, average triangles per edge, and maximum triangle participation. Furthermore, hierarchical organization was evaluated using k-core decomposition and the maximum core number. Community structure was assessed using weighted Louvain detection by counting the number of communities, and finally, global integration was summarized using the network diameter.

For each subject in the held-out test set, all 15 graph statistics were computed on the real and reconstructed networks. The reconstruction accuracy was evaluated using multiple error metrics, including mean absolute error (MAE), root mean squared error (RMSE), relative MAE (RMAE), normalized error, and normalized RMSE (NRMSE). Table 5 illustrates the mean and standard deviation of these metrics across 10 random splits for two independent UK Biobank subsets (UKB1 and UKB2), conditioned on either sex or fluid intelligence. Across all datasets and conditioning variables, the VAE achieves consistently low normalized reconstruction errors, with NRMSE values ranging between 0.148 and 0.156 and normalized error values below 0.08. Both UKB1 and UKB2 indicate strong cross-dataset generalization due to the similarity of error magnitudes between the datasets. The comparable performance across conditioning variables also indicates that the model remains robust to different forms of phenotypic conditioning. Additionally, the relatively small standard deviations across splits demonstrate stable reconstruction of graph statistics and low sensitivity to training variability.

**Table 5:**
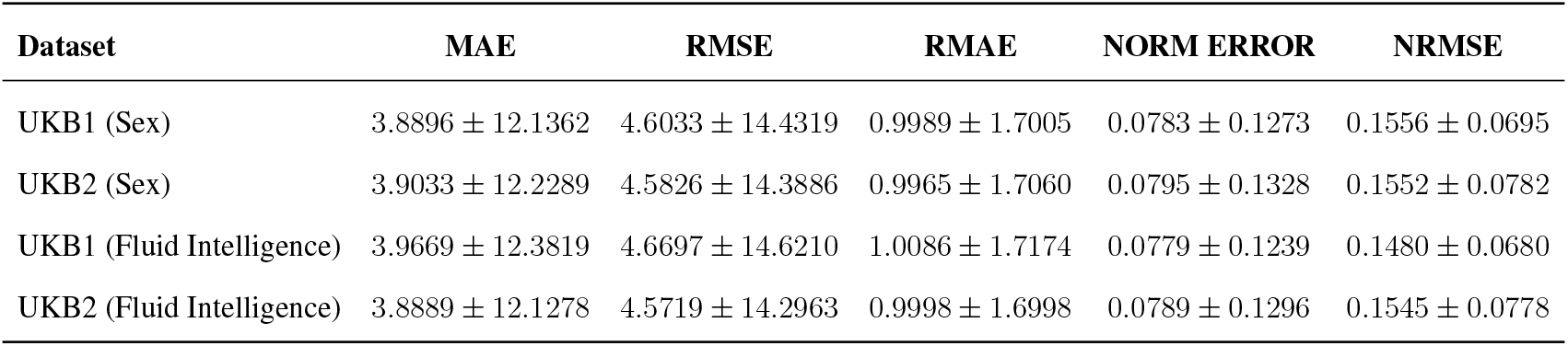
Mean *±* standard deviation of C-GVAE error metrics across UKB datasets and conditioning variables.

| Dataset | MAE | RMSE | RMAE | NORM ERROR | NRMSE |
| --- | --- | --- | --- | --- | --- |
| UKB1 (Sex) | $3.8896 \pm 12.1362$ | $4.6033 \pm 14.4319$ | $0.9989 \pm 1.7005$ | $0.0783 \pm 0.1273$ | $0.1556 \pm 0.0695$ |
| UKB2 (Sex) | $3.9033 \pm 12.2289$ | $4.5826 \pm 14.3886$ | $0.9965 \pm 1.7060$ | $0.0795 \pm 0.1328$ | $0.1552 \pm 0.0782$ |
| UKB1 (Fluid Intelligence) | $3.9669 \pm 12.3819$ | $4.6697 \pm 14.6210$ | $1.0086 \pm 1.7174$ | $0.0779 \pm 0.1239$ | $0.1480 \pm 0.0680$ |
| UKB2 (Fluid Intelligence) | $3.8889 \pm 12.1278$ | $4.5719 \pm 14.2963$ | $0.9998 \pm 1.6998$ | $0.0789 \pm 0.1296$ | $0.1545 \pm 0.0778$ |

To further evaluate reconstruction fidelity across individual graph statistics, the NRMSE was computed for 15 network properties on reconstructed sFNC graphs. Figure. 4 shows a heatmap of the mean NRMSE values per statistic for the four cases: UKB1 and UKB2, conditioned on sex and fluid intelligence. Here, the reconstruction errors are low across all statistics, with minor variation between datasets and conditioning variables. Certain statistics, such as maximum k-core and minimum strength, exhibit slightly higher NRMSE values, reflecting their sensitivity to global or extreme local graph properties. Overall, the heatmap demonstrates that the C-GVAE model consistently preserves the topological structure of functional brain networks across different datasets and phenotypic conditions.

**Figure 4:**
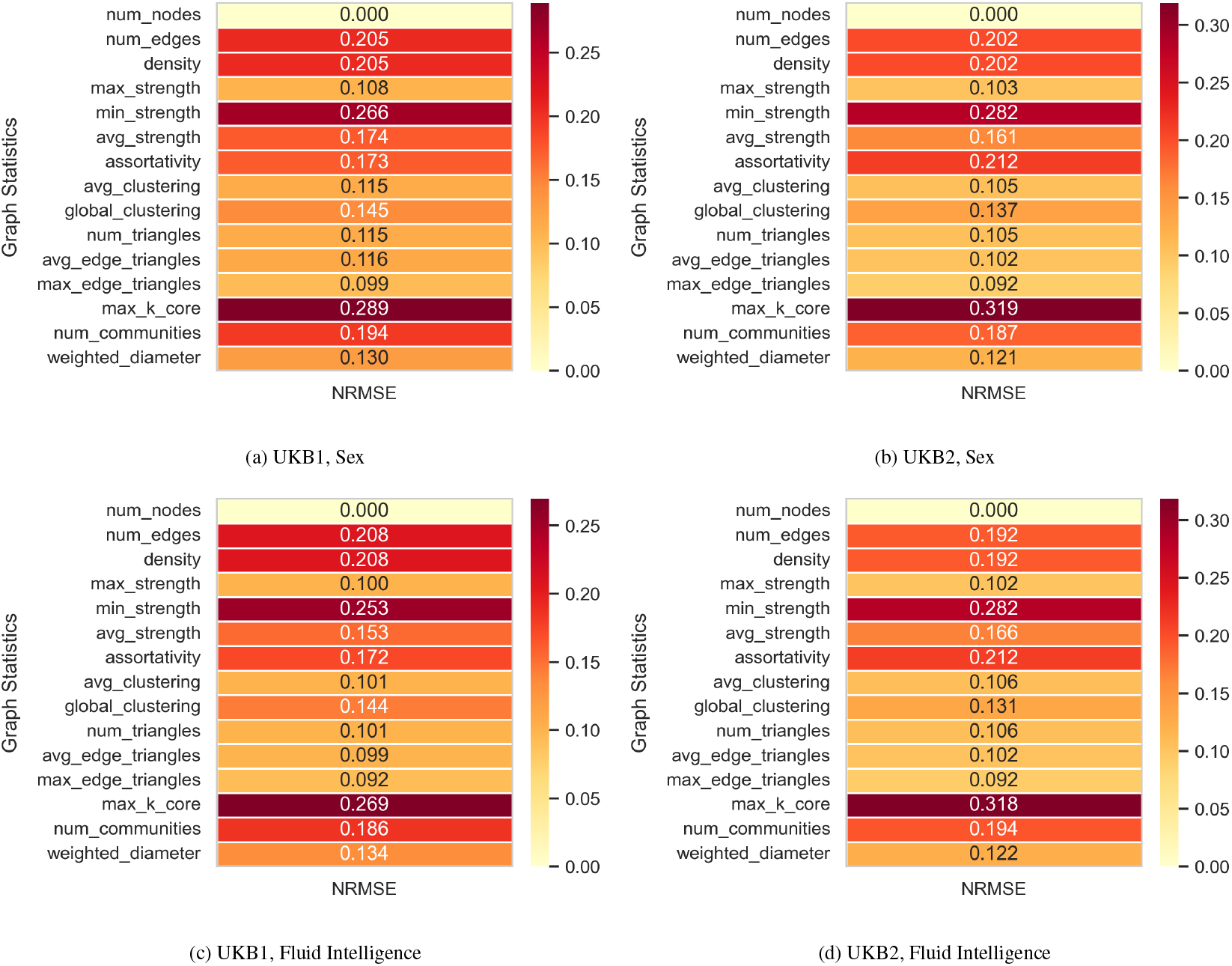
Heatmaps of mean normalized root mean squared error for 15 graph statistics across reconstructed sFNC graphs. Panels show results for UKB1 and UKB2, conditioned on sex (a, b) and fluid intelligence (c, d). Low NRMSE values indicate that the C-GVAE accurately preserves both local and global network properties.

#### Studying the latent representations of real data

To extract the latent vectors for the test set, the trained C-GVAE is utilized without further updating its parameters. From the fixed external test sets of UKB1 and UKB2, samples corresponding to a specific sex are filtered to form a sex-specific subset. Each sample in this subset is then passed through the encoder of the C-GVAE, which incorporates the graph-structured input. The encoder produces intermediate latent embeddings that are concatenated with sex information and passed through linear layers to compute the mean and variance of the latent distribution. Using the reparameterization trick, the latent vectors are sampled from this distribution. These latent vectors are collected for all samples in the test set. We then perform a two-sample t-test with FDR correction for group comparison between the male and female samples in the test sets. Figure 5 shows the results from the group comparison done independently for UKB1 and UKB2 datasets. It can be seen that a significant percentage of latent vectors encode significant group differences between male and female subjects. These elements were selected for further unit-testing of the latent space of the model as described in the subsequent subsection.

**Figure 5:**
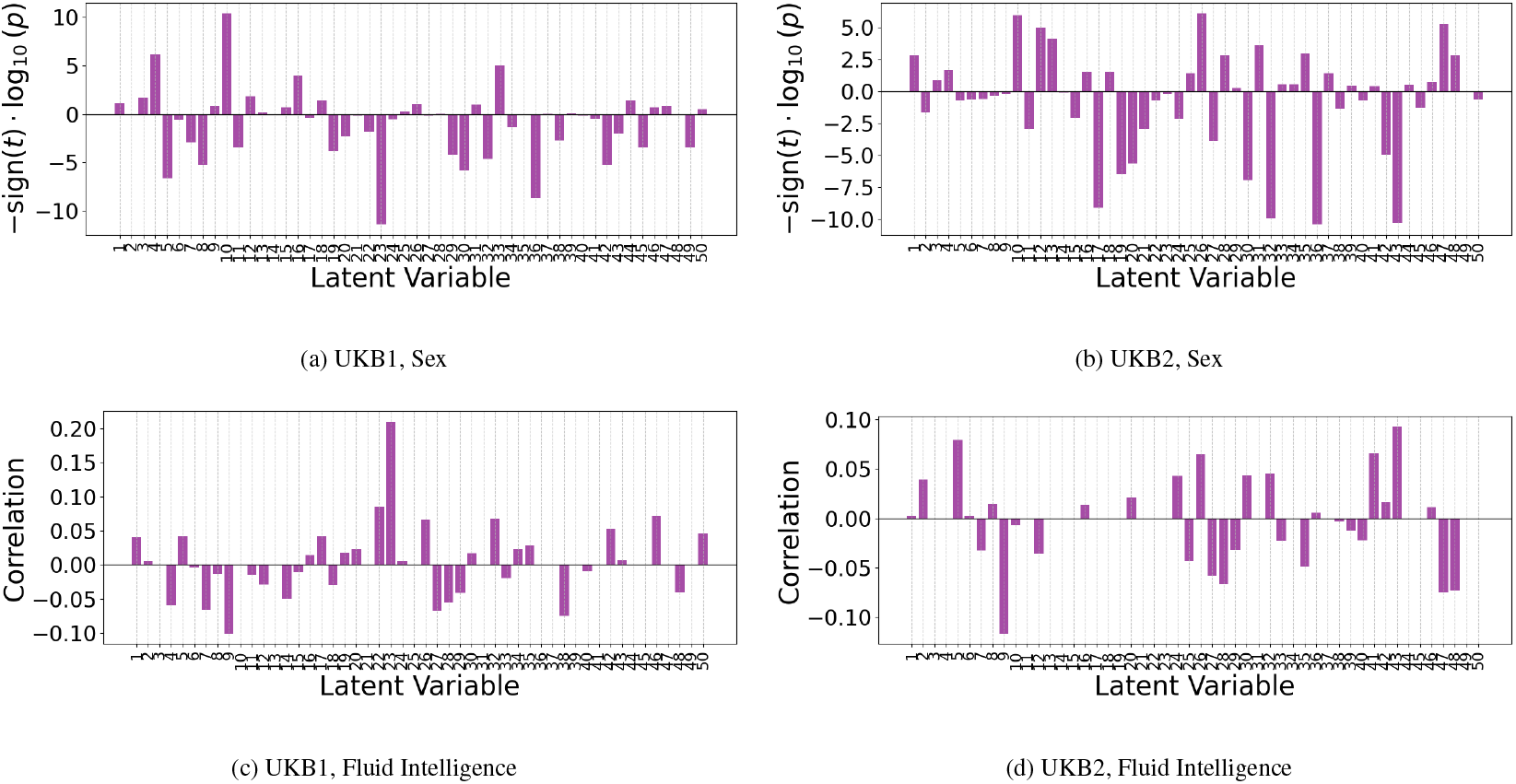
Group differences (−*sign*(*t*) *log*(*p*) values from two-sample t-test) and correlations after FDR correction between the encoded latent vector embeddings from sex and FI-score conditioning, respectively, on the test data of UKB1 and UKB2.

In the case of fluid intelligence, the same test sets were passed through the encoder into the latent layers, but with fluid intelligence score as the a contiuous conditioning variable. In this case, instead of a two-sample t-test unlike sex, we performed correaltion analysis between the encoded latent vector elements and fluid intelligence scores Figure 5.

#### Analyzing the latent space of the C-GVAE model

Latent space analysis was conducted on the trained C-GVAE model, with conditioning based on the sex. After training, the model’s decoder is analyzed systematically to understand how individual latent dimensions influence the reconstructed sFNC output. This is done by performing one-hot probing, where each latent dimension is independently activated (set to 1.0) while others are held at zero, and the decoder reconstructs an adjacency matrix given this latent code and a fixed gender condition (e.g., male or female). The resulting matrices are visualized as heatmaps, showing the functional patterns the decoder associates with activation of each latent feature. This approach provides insight into the generative factors learned by the model underlying the graph data and facilitates comparison across values of the conditioning variable (sex and fluid intelligence score). For this comparison, we looked at the difference between the decoded outputs of male and female conditioning in the case of sex, and high and low values in the case of fluid intelligence scores thresholded at 0 and 1, for each dimension independently. These differences in the decoded outputs for the one-hot representation of latent dimensions are visualized as connectograms (see Figure 6) for the top two dimensions with the most significant differences in the latent vector representation described in section. Essentially, these connectograms represent the functional connections that encode the highest difference based on the conditioning variable in the latent space.

**Figure 6:**
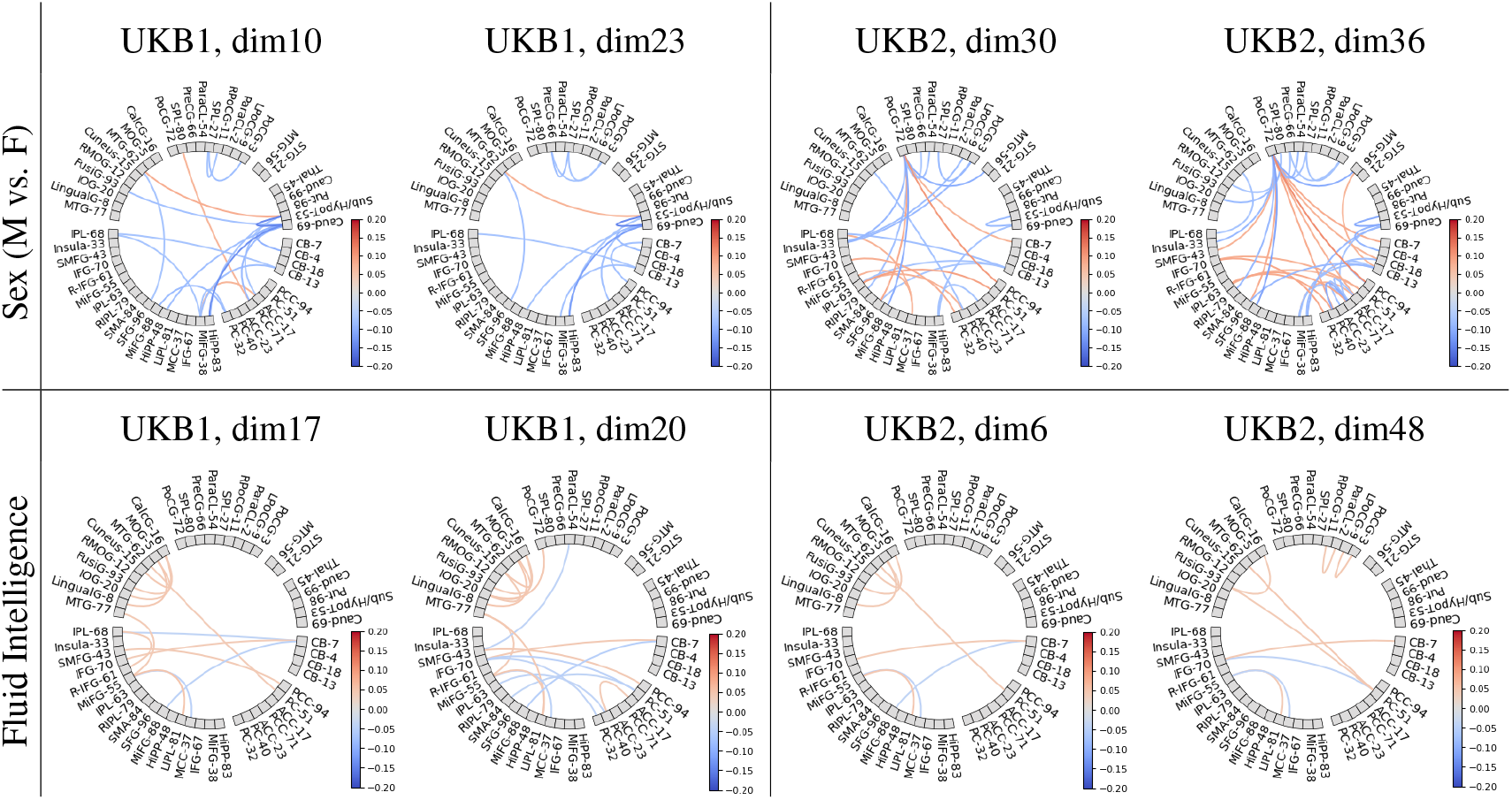
Unit testing of the most discriminative latent vectors of GraphVAE. One-hot encoded values of the two best latent vector elements based on the strongest M vs F (sex) and high vs low (fluid intelligence) group differences for the two conditioning variables were decoded into the sFNC domain as functional connections and are visualized as connectograms above.

## DISCUSSION

Our experimental results demonstrate that the proposed C-GVAE model not only generates realistic, high-quality sFNC data but also captures biologically meaningful patterns. Notably, the group-level differences observed between males and females in the reconstructed data align with established findings on sex-specific functional connectivity. Two-sample t-tests reveal that the reconstructed sFNC data retain key sex-related and fluid intelligence patterns seen in real data. Specifically, the observed sex differences in sFNC features, such as stronger positive correlations within default mode network (DMN) regions and negative correlations in cognitive control areas. Previous studies have shown that females exhibit greater within-DMN connectivity, while males exhibit reduced DMN activation and relatively stronger connectivity in task-positive networks such as cognitive control or frontoparietal networks Ficek-Tani et al. (2023); S. Zhang et al. (2025). Similarly, for fluid intelligence, we observed negative correlations within the cognitive control network, positive correlations within the subcortical domain, and negative correlations between the sensorimotor and cognitive control domains. These findings are consistent with previous studies highlighting the role of subcortical structures, such as the thalamus and basal ganglia, in supporting higher-order cognitive functions Fama and Sullivan (2015).

Additionally, unit testing of the most discriminative latent vectors in GraphVAE revealed how the model captured distinct brain connectivity patterns associated with sex differences and varying levels of fluid intelligence. By one-hot probing the latent dimensions and decoding them back into the sFNC space, distinct connectograms were generated for the most discriminative latent dimensions. This highlighted the strongest functional differences across groups that could be summarized at the level of functional network domains as well. For the male versus female comparisons, strong negative correlations were observed within the subcortical domain, particularly between the caudate and subthalamus/hypothalamus, as well as between the caudate and the hippocampus (cognitive control domain). These findings indicated that sex-related differences in brain organization may be reflected in network domains, which are known to play key roles in memory, learning, and regulatory processes.

In terms of fluid intelligence, the decoded connectograms revealed strong positive correlations between visual domain ICs such as the middle temporal gyrus and the precuneus of the default mode network. Additionally, stronger negative correlations were found between the middle frontal gyrus (cognitive control domain) and the cerebellum, alongside positive correlations between the inferior frontal gyrus (cognitive control domain) and the cerebellum. These connectivity patterns align with the literature, indicating that higher cognitive performance is supported by integrated visual, default mode, and cognitive control network activity, as well as by dynamic interactions between frontal regions and the cerebellum Anat et al. (2024); Thapaliya et al. (2025).

Overall, these results demonstrate that the proposed C-GVAE framework learns structured, biologically grounded representations that preserve meaningful group-level differences across different cognitive and demographic variables. The consistency between reconstructed sFNC patterns, discriminative latent dimensions, and well-established neurobiological findings underscores the interpretability, generalizability, and robustness of the learned representations. Importantly, the identification of group differences at both the level of individual connections and broader functional network domains shows that the model captures brain organization across multiple spatial scales. This makes the proposed C-GVAE model a promising tool for studying inter-subject variability in brain connectivity in a data-driven manner.

## CONCLUSION

By combining attention-based GNNs with conditional variational autoencoders, we present a methodological framework that integrates generative modeling of functional network connectivity for the human brain with learning latent representations that capture characteristic differences associated with clinically observed variables at hand. Our model, trained independently on two sub-cohorts of a large fMRI dataset, generates realistic functional connectivity graphs that retain key network metrics and the essential discretionary characteristics of a given clinically observed variable (biological sex or fluid intelligence score) at hand. Through multiple rigorous statistical analyses, we study these differences in both the latent as well as the reconstructed space of the model. We show that predictive classification performance on unseen data is better when using the reconstructed data, thus enhancing the separability based on the target variable. We also visualize the latent representations, highlighting condition-specific differences within meaningful functional networks that correspond to brain components.

For a successful data-driven development of biomarkers for functional brain disorders, it is essential to create frameworks that utilize the power of generative models to capture the complexities of neuroimaging data. These frameworks should learn meaningful latent representations that map features to one another, while also providing biologically interpretable visualizations of key patterns associated with clinically observed assessments. This work is aimed at achieving this goal. In the future, this work can be extended to include datasets from neurological disorders so as to study more disorder-specific clinical assessments. Another line of extension could be to include multiple modalities so as to study the interplay of behavioral assessments with the structural and functional features of the brain.

## ACKNOWLEDGMENTS

Research reported in this publication was supported by the National Institutes of Health (NIH) under award numbers 1R01AG090597 and R01AG073949.

## AUTHOR CONTRIBUTIONS

IB: conceptualization, formal analysis, methodology design, visualization, writing, reviewing, editing; MA: methodology design, formal analysis, implementation, writing, reviewing, editing; VDC: conceptualization, supervision, administration, resources, reviewing, editing.

## DATA AVAILABILITY

The datasets analyzed during the current study are available from the UK Biobank (https://www.ukbiobank.ac.uk/) and the Human Connectome Project (HCP) (https://www.humanconnectome.org/). These data are not publicly available but can be accessed upon request through the respective data providers.

